# Eye morphogenesis in the blind Mexican cavefish

**DOI:** 10.1101/698035

**Authors:** Lucie Devos, François Agnès, Joanne Edouard, Victor Simon, Laurent Legendre, Naima El Khallouki, Sosthène Barbachou, Frédéric Sohm, Sylvie Rétaux

## Abstract

The morphogenesis of the vertebrate eye consists of a complex choreography of cell movements, tightly coupled to axial regionalization and cell type specification processes. Disturbances in these events can lead to developmental defects and blindness. Here, we have deciphered the sequence of defective events leading to coloboma in the embryonic eye of the blind cavefish of the species *Astyanax mexicanus*. Using comparative live imaging on targeted enhancer-trap *Zic1:hsp70:GFP* reporter lines of both the normal, river-dwelling morph and the cave morph of the species, we identified defects in migratory cell behaviors during evagination which participate in the reduced optic vesicle size in cavefish, without proliferation defect. Further, impaired optic cup invagination shifts the relative position of the lens and contributes to coloboma in cavefish. Based on these results, we propose a developmental scenario to explain the cavefish phenotype and discuss developmental constraints to morphological evolution. The cavefish eye appears as an outstanding natural mutant model to study molecular and cellular processes involved in optic region morphogenesis.

## Introduction

The morphogenesis of the vertebrate eye follows a complex choreography of cell movements, starting from a flat neural plate to generate a spherical multi-layered structure. This process is advantageously investigated on teleost models, which are amenable to live imaging (reviewed in (Cavodeassi, 2018).

At the end of gastrulation, the “eyefield” is specified in the anterior neural plate, surrounded anteriorly and laterally by the prospective telencephalon, and posteriorly by the future hypothalamus and diencephalon (Varga et al., 1999; Woo and Fraser, 1995; Woo et al., 1995). The first step of eye formation is the lateral evagination of the optic vesicles (OV) (England et al., 2006; Ivanovitch et al., 2013; Rembold et al., 2006). The vesicles then elongate due to a flow of cells entering the anterior/nasal OV, in a process recently re-described as “extended evagination” (Kwan et al., 2012). Simultaneously, the OVs are separated from the neural keel by the anterior-wards progression of a posterior furrow (England et al., 2006). Cells from the inner OV leaflet then migrate around the rim of the eye ventricle, the optic recess, into the lens facing neuroepithelium through the “rim movement” (Heermann et al., 2015; Kwan et al., 2012). The cells fated to the retinal pigmented epithelium (RPE) expand and flatten to cover the back of the retina (Cechmanek and McFarlane, 2017; Heermann et al., 2015). Together with the basal constriction of lens-facing epithelial cells (Martinez-Morales et al., 2009; Nicolas-Perez et al., 2016), these movements lead to optic cup (OC) invagination and also to the formation of the optic fissure - which needs to close to have a functional, round eye (Gestri et al., 2018). Finally, the entire eye, together with the forebrain, rotates anteriorly, bringing the fissure in its final ventral position. Hence, cells that are initially located in the dorsal or ventral part of the OV contribute to the nasal or temporal quadrant of the retina, respectively (Picker et al., 2009) (**Fig.S1**). Failure to complete correctly any of these steps can lead to vision defects; for example, failure to close properly the optic fissure is termed coloboma.

*Astyanax mexicanus* is a teleost that arises in two morphs: classical river-dwelling eyed morphs and blind cave-dwelling morphs. Although eyes are absent in adult cavefish, they first develop in embryos before degenerating during larval stages. The embryonic cavefish eyes display several abnormalities: the OVs are short (Alunni et al., 2007), the OC and lens are small (Hinaux et al., 2015; Hinaux et al., 2016; Yamamoto and Jeffery, 2000) and the ventral OC is severely reduced or lacking, leaving the fissure wide open with a coloboma phenotype (Pottin et al., 2011; Yamamoto et al., 2004). Cavefish exhibit several modifications of morphogen expression which trigger changes of the cavefish eyefield and subsequent eye, and which have been linked to cavefish eye defects. Accordingly, overexpression of *Shh* in surface fish shortens its optic cups and triggers lens apoptosis, while inhibition of Fgf signalling in cavefish restores the ventral retina (Hinaux et al., 2016; Pottin et al., 2011; Torres-Paz et al., 2019; Yamamoto et al., 2004) .

Because of these variations, the cavefish is a remarkable natural mutant model to study eye development, beyond the mechanisms of eye degeneration and loss. Here, we sought better understanding cavefish embryonic eye defects as well as the mechanisms of eye morphogenesis in general. We generated CRISPR/Cas9-mediated targeted enhancer trap cavefish and surface fish *Zic1:hsp70:GFP* lines and performed comparative live imaging of eye morphogenesis in developing embryos of the two morphs to uncover the morphogenetic processes and cellular behaviors leading to cavefish coloboma.

## Results and Discussion

### Establishing *Zic1:hsp70:GFP* surface fish and cavefish knock-in reporter lines

We performed an *in situ* hybridization mini-screen to choose a candidate reporter gene labelling the entire optic region from neural plate stage (10hpf) until at least 30hpf (**Fig. S2A**). *Zic1* was chosen due to its early and persistent expression in the optic field (**Figure 1A**; **Fig. S2B** and legend), even though its pattern was complex and larger than the optic region.

**Figure 1:**
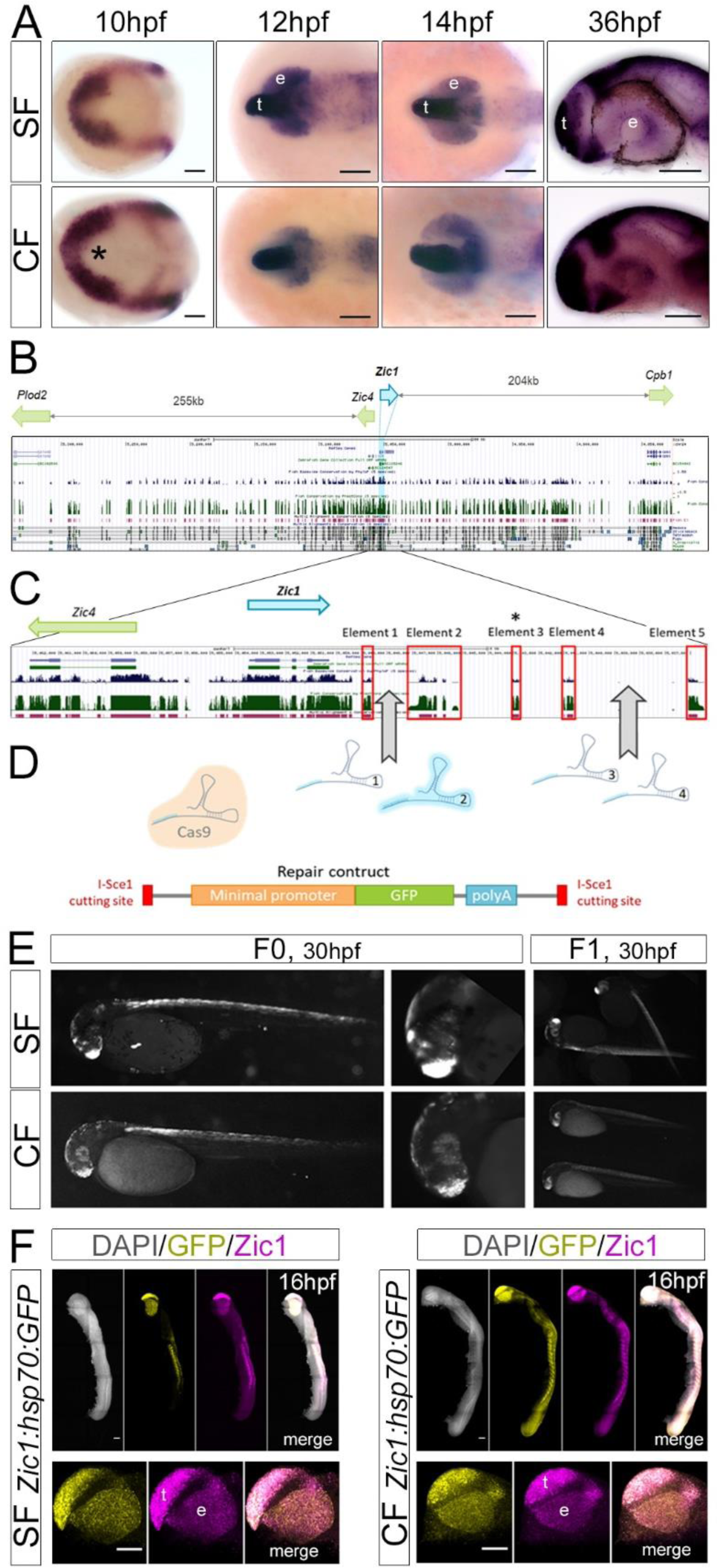
establishment of surface fish and cavefish *Zic1:hsp70:GFP* lines. (A) *Zic1* expression at indicated stages in SF and CF. Anterior is to the left. Dorsal views at 10, 12 and 14hpf, lateral views at 36hpf. Asterisk: larger indentation in the CF eyefield. (B) Zebrafish *Zic1* genomic region in UCSC genome browser (2010 assembly). Green blue peaks as well as magenta and black elements correspond to high conservation, showing the complexity of the region. (C) Close-up on *Zic1*. Red boxes highlight conserved elements; element 3 is not conserved in *Astyanax* (asterisk). (C) sgRNA were designed to target the low-conservation regions between elements 1 and 2, and 3 and 4. SgRNA2 (pale blue) efficiently generated cuts. It was co-injected together with the Cas9 protein and the linear repair construct containing a minimal Hsp70 promoter and the GFP. (D) *Zic1-like* GFP fluorescence in mosaic F0s and stable F1s. (E) Double-fluorescent *in situ* hybridization at 16hpf for *Zic1* (magenta) and *GFP* (yellow) showing that the transgene recapitulates the endogenous *Zic1* pattern, both for SF and CF lines. The top panels show entire embryos and the bottom panels show close-ups on the head, including the *Zic1*-expressing telencephalon (t) and eye (e). Lateral views. Scale bars=100µm.

We used a targeted enhancer-trap strategy into the *Zic1* locus, so that the GFP reporter insertion site would be similar in CF and SF lines and avoid positional effects, which is crucial for comparative purposes. The large and complex *Zic1* genomic region was examined to find conserved elements pointing toward putative regulatory elements (**Fig. 1BC**). In both zebrafish and *Astyanax* genomes (McGaugh et al., 2014), *Zic1* and *Zic4* were located in a head to head configuration in the middle of a gene desert (∼275kb downstream of *Zic1* and ∼235kb downstream of *Zic4* in *Astyanax)* which contained many fish-conserved elements, also partly conserved with tetrapods (**Fig. 1BC**). Such a regulatory landscape suggested that the elements driving *Zic1* expression are probably modular and difficult to identify, further strengthening the choice of a directed enhancer-trap approach. We thus “addressed” the enhancer-trap construct to *Zic1* downstream region using CRISPR/Cas9, similarly to the approach used by Kimura and colleagues (Kimura et al., 2014). We reasoned that using NHEJ (non-homologous end joining) DNA repair mechanism-based strategy, the preferred repair mechanism in fish embryos (Hagmann et al., 1998), would maximize integration efficiency. CF and SF eggs were co-injected with sgRNA2 (targeting the region between conserved non-coding elements 1 and 2), Cas9 protein and a linearized minimal promoter *hsp70:GFP* repair construct, and embryos were screened at 30hpf for fluorescence patterns consistent with *Zic1* endogenous expression (**Fig. 1E**). This method yielded good results, as its limited efficiency was compensated by the possibility of using a pattern-based fluorescence screening in F0 embryos. Excellent *Zic1* pattern recapitulation in F0 was observed at low frequency (1-2% of injected embryos), while other, more partial patterns were seen at higher frequencies. All potential founder embryos were raised until males were sexually mature (6 months) and could be screened by individual *in vitro* fertilization. We detected 3 founders for SF (out of 15 F0 males screened) and 5 founders for CF (out of 9 F0 males screened) with various transmission rates: 4%, 7% and 30% for SF founders and 4%, 45%, 48%, 50% and 54% for CF founders, respectively. Hence, we obtained an excellent ratio of founder fish among selected F0 embryos (>50% in cavefish). The fish were screened based on their GFP pattern, matching *Zic1* (**Fig. 1E**). In both morphs some variations in relative fluorescence intensities were observed, with some lines exhibiting homogeneous expression levels and others showing strong GFP fluorescence in the telencephalon and dimer fluorescence in the eye. We focused on the most homogeneous lines for imaging purposes. Importantly, in those lines, genomic analyses confirmed the proper insertion of the transgene at the targeted site, although some structural differences existed (**Fig. S3**). The insertion method being based upon non-conservative NHEJ mechanism, these variations are likely due to sequence differences from one line to another (indels or duplications in genomic DNA or transgene), which may affect the nearby regulatory sequences and slightly modify transgene expression. However, such variations remain anecdotal compared to the differences observed between lines generated by traditional transgenesis techniques (such as Tol2 transgenesis) (Elipot et al., 2014; Hinaux et al., 2015; Stahl et al., 2019), validating this approach as a valuable tool to follow gene expression in *Astyanax* morphotypes. Finally, double fluorescent *in situ* hybridisation for *Zic1* and *GFP* mRNAs demonstrated that the reporter fully recapitulated the endogenous *Zic1* pattern at the stages of interest (**Fig. 1F**).

CRISPR/Cas9 has been reported in surface *Astyanax mexicanus* to generate an *Oca2* null mutant and to confirm the role of *Oca2* in the control of pigmentation (Klaassen et al., 2018). This is to our knowledge the first report of the CRISPR/Cas9 technology used in this emergent model species to generate identical reporter lines in the two morphotypes, and in a targeted genome edition perspective.

### Comparing eye morphogenesis in surface fish and cavefish through live imaging

Live imaging was performed on a light-sheet microscope on *Zic1:hsp70:GFP* lines from ∼10.5hpf to 24-30hpf (**Fig. 2** and **Movies 1 and 2**). Embryos were injected with H2B-mCherry mRNA to follow cell nuclei. The orthogonal illumination of the SPIM induced minimal photo-damage, and embryos developing for more than 20hours under the microscope were alive with a normal head shape at 48-60hpf -even though the tail was usually twisted due to the mechanical constraint in the low-melting agarose.

**Figure 2:**
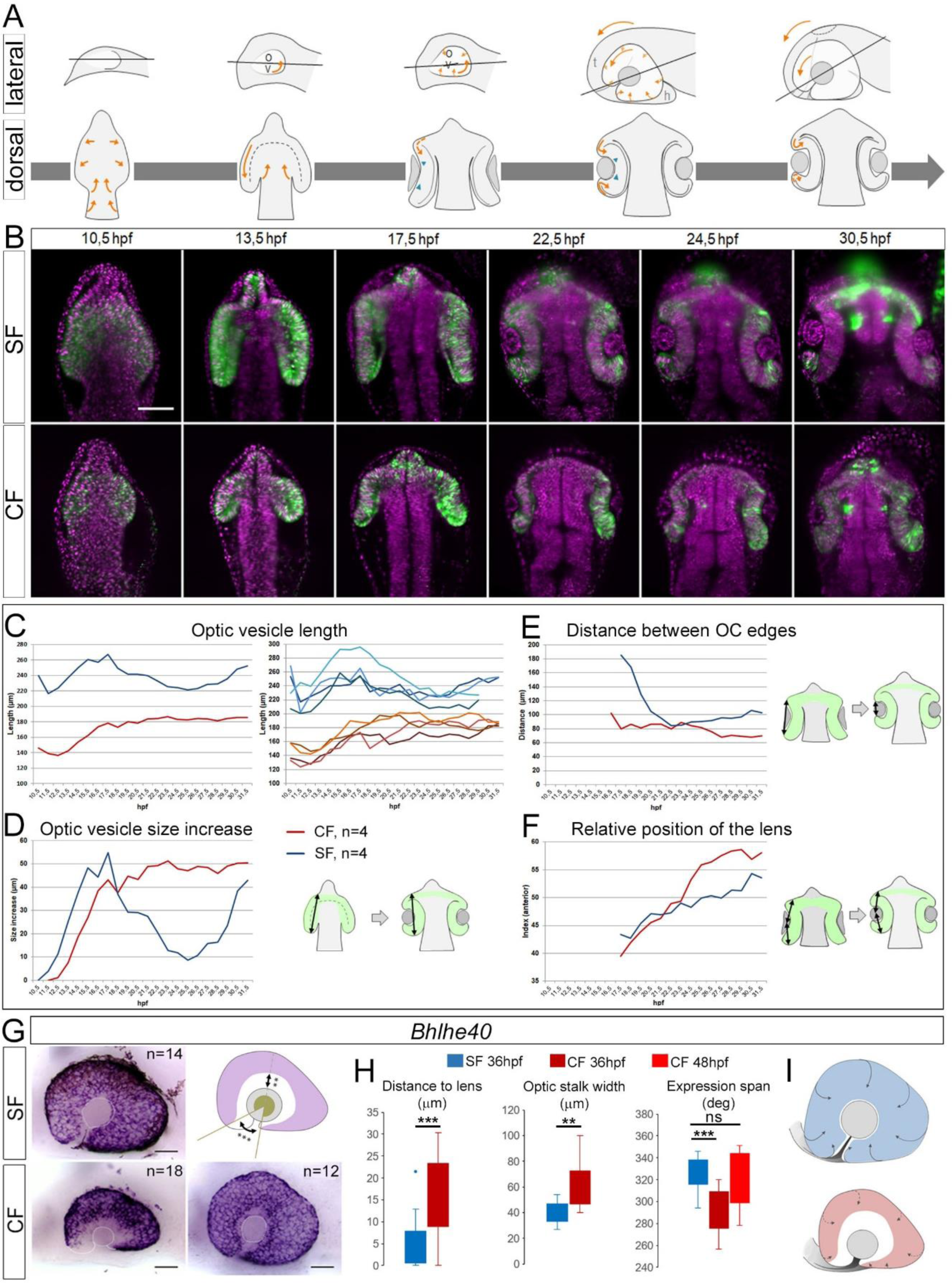
Live imaging and quantification of surface fish and cavefish eye morphogenesis. (A) Schematic drawings of the main steps of eye morphogenesis in fish, in lateral (top) and dorsal views (bottom). Orange arrows indicate cell and tissue movements; green arrowheads show initiation of basal constriction. The grey line indicates the optical section plane used in the pictures in B, which follows an optic stalk to lens center axis and accompanies the anterior rotation illustrated by the arrows. All measures in C-F were done on these planes. (B) Still images of time-lapse acquisitions from 10.5hpf to 30.5hpf on SF (top) and CF (bottom) *Zic1:hsp70:GFP* lines (green, GFP; magenta, nuclear mCherry). Representative steps of eye morphogenesis illustrating CF/SF differences are shown. Dorsal views, anterior to the top. (C-F) Measurements. (C) OV length. The left graph shows the mean of n=4 eyes in each morph (blue, SF; red, CF); the right graph displays the trajectories of individual eyes, showing the reproducibility of the results. Measures are illustrated on the diagrams on the right. (D) OV size increase. (E) Distance between the two optic cup edges. (F) Position of the lens relative to anterior OV, showing that the lens is progressively shifted anteriorly between 25hpf and 30hpf. (G-I) *Bhlhe40* expression. (G) In situ hybridization at 36hpf (left) and 48hpf (bottom right, CF). The scheme shows the measures taken in (H). (I) Drawings illustrating comparative RPE spreading in SF (top) and CF (bottom). Mann-Whitney test: ** p<0.01; *** p<0.001.

For analysis, we chose a plane crossing the middle of the lens and the optic stalk (lines on **Fig. 2A**), to follow the anterior rotation of the eye. Overall, optic morphogenesis in SF recapitulated the events described in zebrafish, while in CF the movements were conserved but their relative timing and extent appeared different. The following macroscopic analyses result from quantifications made on n=4 eyes for each morph.

#### Evagination and elongation of the OVs

The CF OVs were about half-shorter than the SF OVs from the beginning of evagination onwards (139µm *vs* 216µm at 11.5hpf) (**Fig. 2A-C**). Elongation progressed at about the same pace as in SF until 17.5hpf (**Fig. 2C**). However, while OV length decreased between 17.5-25.5hpf in SF due invagination, elongation continued at slower pace until 25.5hpf in CF (**Fig. 2CD**). Moreover, the final size of the SF OC was very similar to the early evaginating eyefield (240µm at 10.5hpf *vs* 252µm at 31.5hpf) while in CF a net increase was observed (146µm at 10.5hpf *vs* 186µm at 31.5hpf) (**Fig. 2C**). In addition, in SF the OVs remained closely apposed to the neural tube, while in CF they first started growing away before getting back closer between 18.5-21.5hpf (**Fig. 2B**). Finally, throughout development, the width of the optic stalk (defined in its wide meaning as the connection between OVs and neural tube) was similar in the two morphs (**Fig. S4**), despite an initially smaller size in CF due to the smaller OVs.

Since elongation proceeds at a similar rate in CF and SF until 17.5hpf, the shorter size of the cavefish OV (Alunni et al., 2007; Strickler et al., 2001) seems principally due to the small size of the initial eyefield (Agnès et al., 2021). Of note, albeit smaller, CF OVs seem “correctly” patterned in their future naso-temporal axis, according to *FoxG1* and *FoxD1* markers at 13.5hpf (Hernandez-Bejarano et al., 2015). Then, after the initial evagination and patterning of small OVs, morphogenesis proceeds with the extended evagination, whereby cells from the neural tube continue entering the OV to contribute exclusively to the ventro-nasal part of the eye (Kwan et al., 2012). Our measurements suggest that this step proceeds normally in CF. This could partially compensate the originally small size of the eyefield/OV, but only in the nasal part, while the temporal part would remain affected in size.

#### Optic cup invagination and lens formation

The posterior end of the OVs started curling back in both CF and SF around 15.5hpf. The lens was identifiable as an ectodermal thickening at 17.5hpf (**Fig. 2B** and **movies 1 and 2**), in a central position with regard to the antero-posterior extension of the OV, in both morphs (**Fig. 2B,F**). Then, in SF, invagination quickly brought closer the two OC edges in contact with the lens (**Fig. 2B,E**). In contrast, despite initially harbouring a curvature typical of invagination, in CF the OC edges remained flat, with an apparent impairment of the rim movement in their posterior part (**Fig. 2B,E and Movie 2**). The CF OVs continued to elongate while the lens remained static, therefore shifting the lens position anteriorly (**Fig. 2B,F**). The posterior OC showed slow and reduced curling, which in some cases led to a separation from the lens. Eventually, the posterior (prospective dorsal) OC finally curved and contacted the lens (**Movie 2**; **Fig. 2B**), but remained shallower with small bulging lens.

Thus, although the invagination in CF seems to start normally between 15.5-19.5hpf, it progresses poorly so that the OCs remain elongated. This timing is reminiscent of the two steps described for OC invagination in zebrafish: basal constriction initiates the primary folding between 18-20hpf (18-22ss), followed by the rim movement which brings the presumptive retina from the inner OV leaflet into the lens-facing epithelium by an active migration around the rims of the optic recess between 20-24hpf (Heermann et al., 2015; Nicolas-Perez et al., 2016; Sidhaye and Norden, 2017). In *Astyanax*, 18ss corresponds to ∼16.5hpf (Hinaux et al., 2011), suggesting that the initial basal constriction leading to the onset of OC invagination is conserved in cavefish. In contrast, the prolonged extension and the weak curvature of the OVs suggest that the rim movement must be impaired. We suggest that a continuous flow of cells entering the retina leads to its elongation, in the absence of an efficient rim movement. The later is weaker but not absent in CF, as the posterior OC still manages to contact the lens, but at later stages. Such defective rim movement might be due to various causes, such as defects in the basal membrane or failure to establish proper focal adhesion as seen in the *ojoplano* medaka mutant (Martinez-Morales et al., 2009; Nicolas-Perez et al., 2016; Sidhaye and Norden, 2017). Alternatively, active migration could be altered by extrinsic signals, as in BMP overexpression experiments where the cell flow toward the lens-facing epithelium is reduced (Heermann et al., 2015). The various morphogen modifications known in cavefish, and the fact that the ventral eye can be restored by delaying the onset of Fgf signalling in CF to match the SF timing (Pottin et al., 2011), support this possibility.

It was proposed that spreading and migration of RPE cells is concomitant with the rim movement and may contribute to it as a driving force (Cechmanek and McFarlane, 2017; Moreno-Marmol et al., 2018). In 36hpf SF embryos, the RPE marker *Bhlhe40* was expressed all around the eye, often contacting the lens (**Fig. 2GH**), which we took as an indicator of the correct engulfment of the retina by the migrating RPE. The expression spanned 326° around the eye (**Fig. 2GH**). Conversely, in CF, *Bhlhe40* expression showed a significantly diminished covering of the retina by the RPE (289°), with a wider ventral gap possibly corresponding to wider optic fissure opening and *Bhlhe40*-positive cells further away from the lens, suggesting reduced or delayed retina covering by the RPE (**Fig. 2GHI**). At 48hpf however, the staining span was no longer different from the 36hpf SF. These data show that RPE identity is maintained in the CF eye, yet the expansion and engulfment movement of this tissue to cover the whole retina is delayed compared to SF - reinforcing the idea that the rim movement is impaired in cavefish. Potentially, RPE spreading may also be involved in optic fissure closure, as suggested by the presence of a coloboma upon impairment of the rim movement by *BMP4* overexpression in the OV (Heermann *et al*., 2015). Deficiency in RPE spreading might participate in the cavefish coloboma phenotype (**Fig. 2I**). Interestingly, the transplantation of a healthy SF lens into the CF OC rescues the eye as a structure, i.e., prevents lens-induced degeneration, but does not rescue coloboma (Yamamoto and Jeffery, 2000). This is consistent with our findings showing that improper closure of the fissure is autonomous to CF retinal tissues and results from defective morphogenetic movements.

Finally, our movies show that the lens forms in proper place and time, in both morphs, with regard to initial OC invagination. It is only at later stages that the lens appears more anterior (i.e., facing the presumptive ventral retina after final eye rotation) in cavefish. This apparent displacement of the lens relative to the retina is not due to a movement of the lens itself - which remains fixed throughout eye morphogenesis (Greiling and Clark, 2009), attached to the overlying ectoderm from which it delaminates around 22hpf in *Astyanax* (Hinaux et al., 2017) -but rather to persistent OV elongation. This suggests that proper initial interactions occur between the central OV and the lens to adjust their relative position and to initiate OC invagination. Indeed, in chick, the pre-lens ectoderm is required for OC invagination while the lens placode itself is dispensable (Hyer et al., 2003). In cavefish, such mechanisms could exist and lead to the proper initiation of OC folding, as we have observed. Finally, the anterior-shifted position of the lens, due to elongation without invagination, explains how the lens is ventrally-displaced in the mature CF eye after the final anterior rotation movement, leading to coloboma (**Fig. 2A and I**).

In sum, our live-imaging experiments suggest that, in CF (1) OVs are reduced in size after the initial evagination, (2) OV elongation occurs properly, while (3) invagination is transiently compromised. Below we started addressing the cellular behaviors that may underlie these phenotypes.

### Comparing cell behaviors in surface fish and cavefish during evagination

To study cell behaviors that might contribute to the small size of CF OVs, we tracked cells during evagination, between 11.5hpf-13hpf (1h40, 40 movie frames).

#### OV cells proliferation

Division rates may account for size differences between SF and CF OVs. To test this hypothesis, we reconstructed the complete mitotic pattern of the anterior neural tube or head, in one CF and one SF embryo. Metaphase plates were searched manually and tracked at each time step through the depth of the embryos (**Movies 3-6 and Fig. 3AB**). A total of 1073 and 803 cell divisions were annotated in SF and CF, respectively, during the 100min studied. It is, to our knowledge, the first report providing an estimation of the mitotic rate, ∼10 mitoses per minute in the brain/head, during fish neurulation, and a description of cell mitotic behaviors in the evaginating OVs. In both morphs, mitoses were evenly distributed in time and in space - not considering the strong tendency of mitoses to occur close to ventricles (below and **Fig. S5**). After manual re-segmentation through movie stacks to count mitoses in regions of interest, we found about twice more cell divisions in the SF than in the CF OVs (mean left/right: 154/SF *vs* 67/CF) (**Fig. 3A-E; Fig. S6**). The same was true for the “prospective lens”, i.e. the ectoderm in direct contact with the OVs (17/SF *vs* 7/CF). Such SF/CF difference in the number of mitoses was not observed in a medial neural tube region used as control (157/152)(**Fig. 3DE**). In both OVs and presumptive lens ectoderm, the left/right symmetry of mitoses distributions and numbers was excellent, suggesting that the mitotic landscape was accurately reconstituted. To compare mitotic rates in SF and CF optic tissues, the numbers of mitoses were normalized to OV volumes, in two different ways (**Fig. 3CDE; Fig. S6**). Unexpectedly, the normalized mitotic activity appeared higher in cavefish OVs, suggesting that proliferative activity in the CF optic region somehow tends to compensate for small eyefield size (Agnès et al., 2021), and in any case does not participate in the establishment of OV size differences. Importantly, the mitotic behaviors of SF and CF optic cells were also qualitatively identical. The migration towards the ventricle (optic recess), the orienting/rotating behavior of metaphasic plate cells before dividing, and the post-mitosis integration of daughter cells into the neuroepithelium were systematically observed in both morphs (**Fig. 3F-I**; **Fig. S7**). These results rule out an early proliferative defect in CF OVs to explain their small size, which parallels studies at later stages which dismissed a role for defective proliferation during CF eye degeneration (Alunni et al., 2007; Strickler et al., 2002). The cavefish OVs also appear like an outstanding model to study developmental mechanisms controlling organ size and developmental robustness (Young et al., 2019).

**Figure 3:**
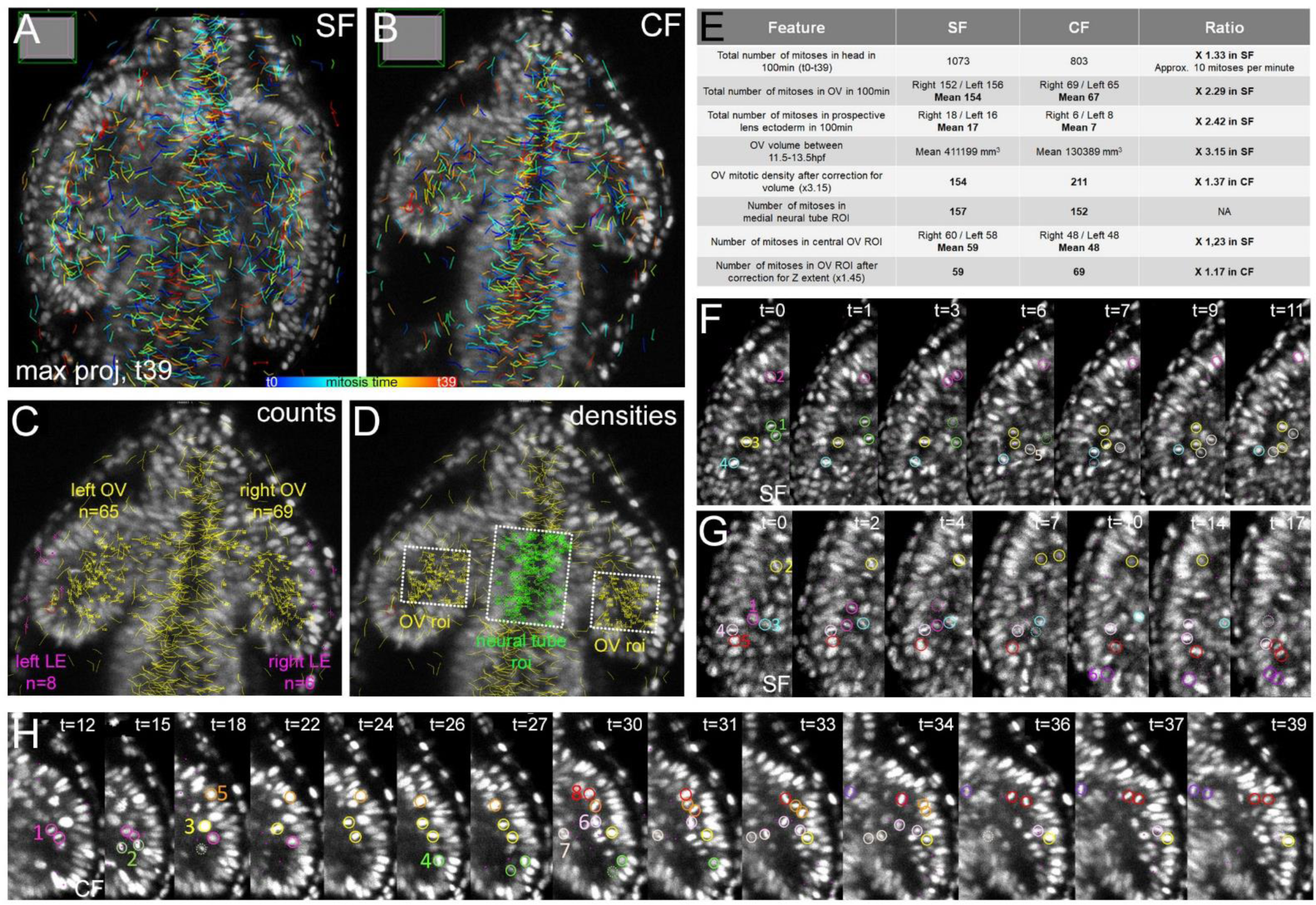
Cell divisions. (A,B) Mitotic embryos. All mitoses tracked during 100 minutes (40 time steps*2.5min) are shown on maximum projection dorsal views at t=39 (end of the movies) in SF (A) and CF (B). The color code indicates division time (see also **Fig. S5**). (C) Mitosis counts, shown here on CF. After completion of tracking, each mitotic event was counted and re-allocated to regions of interest after manual re-segmentation (**Fig. S6**). Mitoses with yellow numbers belong to OVs, while mitoses with pink numbers belong to presumptive lens ectoderm. (D) Mitosis densities were calculated to normalize for OV size differences in the two morphs, shown here on CF. The number of mitoses in a region of interest (roi) of identical size, either at the level of the OVs (yellow numbers) or the medial neural tube (green numbers), were counted in SF and CF (**Fig. S6**). (E) Mitosis quantification and SF/CF comparison. (F,G,H) Cell division behaviors. Colored circles help following individual cells. Representative examples are shown in SF (F,G) and CF (H) OVs (time step: 2.5min). They were qualitatively indistinguishable between SF and CF (more in **Fig. S6**).

#### OV cells trajectories

Defective migratory properties might also contribute to the formation of small OVs in CF. To test this hypothesis, 24 SF and 44 CF OV cells were tracked between 11.5hpf-13hpf (**Fig. 4**).

**Figure 4:**
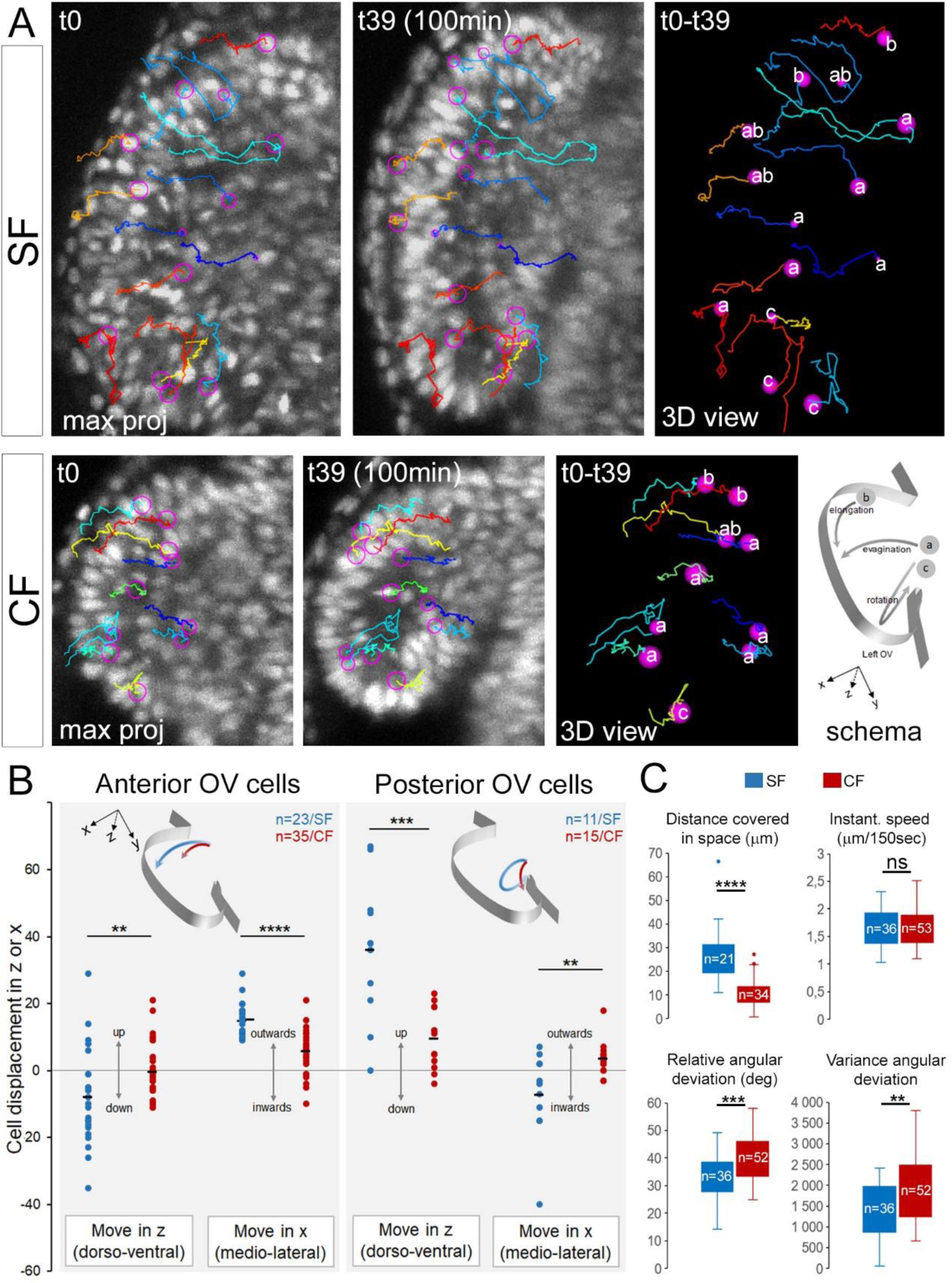
Cell trajectories. (A) Cell tracking and trajectories. Representative examples of cells tracked during 100 minutes, shown on maximum projection dorsal views at t=0 and t=39 (start/end of the movies) and on 3D views at t=0. Individual cell tracks are in different colors, in SF (top) and CF (bottom); cell positions are in pink circles. The bottom right schema illustrates the 3 main types of trajectories (a/evagination; b/elongation; c/rotation). (B) Quantifications of trajectories and directions followed by cells of the 2/3 anterior *versus* 1/3 posterior OV, in SF (blue) and CF (red). (C) Cell migration parameters in SF (blue) and CF (red). Mann-Whitney tests: * p<0.05; ** p<0.01; *** p<0.001.

In SF, we observed markedly different types of trajectories depending on the initial position of cells. Namely, cells located in the 2/3 anterior OV showed a lateral-wards movement with a slight tendency to dive towards the ventral side, thus strongly contributing to evagination (**Fig. 4A-B**). Some anterior cells, either dorsally or ventrally located, also made a posterior turn or had a strict antero-posterior trajectory, potentially contributing to elongation (**Fig. 4A-B**). Conversely, cells located in the posterior third of OVs followed a dorsal-wards and inwards path, seemingly imposing a rotational movement to the posterior OV (**Fig. 4A-B**), and putatively corresponding to the “pinwheel movement” described in zebrafish by (Kwan et al., 2012).

Most of these trajectories were impaired in CF (**Fig. 4A-B**). Anterior cells showed reduced outwards movement and remained static in Z, showing reduced contribution to evagination. Posterior cells trajectories had less amplitude in the upwards direction and displayed outwards instead of inwards trajectories. On the other hand, cells with posterior-wards trajectories contributing to elongation were observed in CF (**Fig. 4A**), in line with the proper elongation recorded above (**Fig. 3**). These data suggested that CF optic cells adopted improper behaviors in terms of trajectories during evagination.

We analysed kinetic parameters of cell migrations. The instantaneous speed and the total distance travelled by OV cells in the two morphs were similar (**Fig. 4C**), suggesting that the migrating apparatuses and capacities of CF cells were unaffected. However, the total displacement in space was markedly shorter for CF cells, in line with above observations on trajectories. To reconcile these apparently contradictory observations, we measured deviation angles of cell trajectories between different time steps. We found a significant zigzagging or erroneous aspect of CF cells migration, as compared to the straighter paths of SF cells (**Fig. 4C**). These data suggest that CF optic cells partly lacked or failed to respond to guidance and directionality cues.

### Conclusions

In all eyeless or eye-reduced cave vertebrates examined so far, initial eye development occurs (e.g., (Durand, 1976; Stemmer et al., 2015; Wilkens, 2001). This represents an energetically-costly process for embryos, raising the puzzling question of why would these species first develop eyes which are after all fated to degeneration, and suggesting that initial eye development cannot be circumvented (Rétaux and Casane, 2013). Our results help refine the step(s) in eye morphogenesis that are subjected to developmental constraint. In cavefish, the eyefield is specified and the evagination/elongation steps, corresponding to cell movements leading to the sorting of retinal versus adjacent telencephalic, preoptic and hypothalamic cells of the neural tube, do occur. It is only after the segregation between these differently-fated cell populations that cavefish eye morphogenesis starts going awry, with a defective invagination process, soon followed by lens apoptosis and progressive degeneration of the entire eye. Therefore, our data support the idea that the first steps of eye development constitute an absolute developmental constraint to morphological evolution. To the best of our knowledge, the closest to a counter-example is the medaka mutant *eyeless*, a temperature-sensitive *rx3* mutant line in which OVs do not evaginate. However, the homozygous *eyeless* fish either die after hatching (Winkler et al., 2000) or, for the 1% which reach adulthood, are sterile probably due to anatomical hypothalamic or hypophysis defects (Ishikawa et al., 2001) -still reinforcing the hypothesis of a strong developmental constraint on vertebrate eye morphogenesis.

Thanks to genome-editing and live-imaging methods, we have started deciphering the morphogenetic and cellular processes underlying colobomatous eye development in cavefish. Further analyses will refine the current scenario. Our data also pave the way for experiments aiming at understanding the defective molecular or signalling mechanisms in cavefish eye morphogenesis, using the *Zic1:hsp70:GFP* knock-in lines and embryology methods we have recently developed (Torres-Paz and Rétaux, 2021).

## Methods

### Animals

Laboratory stocks of *A. mexicanus* surface fish and cavefish were obtained in 2004 from the Jeffery laboratory at the University of Maryland. The surface fish were originally collected from San Solomon Spring, Texas and the cavefish are from the Pachón cave in Mexico. Surface fish are kept at 26°C and cavefish at 22°C. Natural spawns are induced after a cold shock (22°C over weekend) and a return to normal temperature for surface fish; cavefish spawns are induced by raising the temperature to 26°C. Embryos destined for *in situ* hybridization were collected after natural spawning, grown at 24°C and staged according to the developmental staging table (Hinaux et al. 2011) and fixed in 4% paraformaldehyde. After progressive dehydration in methanol, they were stored at -20°C. Embryos destined to transgenesis or live imaging were obtained by *in vitro* fertilization. Embryos were raised in an incubator until 1 month post fertilization for the surface fishes and two month post fertilization for the cavefish. They were kept at low density (15/20 per litre maximum) in embryo medium, in 1 litre plastic tanks with a soft bubbling behind the strainer. Larvae were fed from day 5 with paramecium and transitioned to artemia nauplii from day 10-15. Artemia were given twice a day except for the weekends (once a day) and carefully removed afterward to avoid polluting the medium. At least two thirds of the medium were changed every day and dead larvae removed. After one month for the surface fish and two months for the cavefish, juveniles were taken to the fish facility where they were fed dry pellets (Skretting Gemma wean 0.3) and quickly moved to bigger tanks in order to allow their fast growth.

Animals were treated according to French and European regulations of animals in research. SR’ authorization for use of animals in research is 91-116, and Paris Centre-Sud Ethic committee authorization numbers are 2012-52 and 2012-56.

### *In situ* hybridization

Some cDNAs were available from our cDNA library : *Zic1* (FO290256), *Zic2a* (FO320762) and *Rx3* (FO289986); others were already cloned in the lab : *Lhx2* (EF175737) and *Lhx9* (EF175738) (Alunni et al. 2007); obtained from other labs (*Vax1* : Jeffery lab, University of Maryland; (Yamamoto et al. 2004)); or cloned for the purpose of this work in pGEMT-Easy (Promega) :

- Vax2: forward primer GGGCAAAACATGCGCGTTA; reverse primer CAGTAATCCGGGTCCACTCC.
- Bhlhe40: forward primer : GCACTTTCCCTGCGGATTTC; reverse primer : TGGAGTCTCGTTTGTCCAGC

cDNAs were amplified by PCR, and digoxygenin-labelled riboprobes were synthesized from PCR templates. Embryos were rehydrated by graded series of EtOH/PBS, then for embryos older than 24hpf, proteinase-K permeabilization at 37°C was performed for 36hpf embryos only (10 µg/ml, 15 min) followed by a post-fixation step. Riboprobes were hybridized for 16 hr at 65°C and embryos were incubated with anti-DIG-AP (Roche, dilution 1/4000) overnight at 4°C. Colorimetric detection with BCIP/NBT (Roche) was used. Mounted embryos were imaged on a Nikon Eclipse E800 microscope equipped with a Nikon DXM 1200 camera running under Nikon ACT-1 software. Brightness and contrast were adjusted using FIJI, some of the images used for illustration purpose were created from an image stack, using the extended depth of field function of Photoshop CS5. Area, distance and angle measurements were performed using FIJI (Schindelin et al., 2012).

### In vitro fertilization (IVF) and injections

Surface and cavefish were maintained in a room with shifted photoperiod (light: 4pm – 7am, L:D 15:11) in order to obtain spawns during the working day (*Astyanax* spawn at night (Simon et al., 2019)). Fish activity was monitored after induction and upon visible excitation or when first eggs were found at the bottom of the tank, fish were fished. Females were processed first to obtain eggs: they were quickly blotted on a moist paper towel and laid on their side in a petri dish. They were gently but firmly maintained there while their flank was gently stroked. If eggs were not released immediately, the female was put back in the tank. Once eggs were collected, a male was quickly processed similarly to females, on the lid of the petri dish to collect sperm. The sperm was then washed on the eggs with 10-20mL of tank water (conductivity ∼500µS) and left for a few moments (30s to 2 min approximatively), after which embryo medium was added in the petri dish. Fertilised eggs were quickly laid on a zebrafish injection dish containing agarose grooves. They were injected with a Picospritzer III (Parker Hannifin) pressure injector.

### CRISPR injections and Knock-In lines

sgRNA were designed to target the low-conservation regions between elements 1 and 2 and between elements 3 and 4. Two sgRNA were initially designed per region and sgRNA2 was found to efficiently cut the targeted region (**Fig. S8)**. The mix contained Cas9 protein generously provided by TACGENE and sgRNA2 with the following targeting sequence: CCCAATTCACCAGTATACGT (synthesized with AMBION T7 MEGAshortscript^TM^ T7 transcription kit). Concentrations were kept with a 1:1.5 Cas9 to sgRNA molar ratio and varied between 0.71µM (25ng/µL) and 5.67µM (200ng/µL) of sgRNA 2, mostly 2.84 and 1.42µM were used. The donor construct contained a HSP70 promoter used as a minimal promoter, a GFP cDNA and SV40 poly-adenylation signal, flanked by I-SceI meganuclease cutting sites. I-SceI was used to generate sticky ends and was either detached by 7 min at 96°C or injected with the construct. Concentrations of the repair construct varied between 3.33 and 10.92nM but were mostly used at 10.71nM.

### mRNA injection

Transgenic embryos used for live imaging were injected in the cell or yolk at 1 cell stage with a H2B-mCherry fusion mRNA at a concentration of 50ng/µL.

### Imaging

Transgenic embryos were obtained by IVF with wild-type eggs and transgenic sperm and were immediately injected with H2B-mCherry mRNA for nuclear labelling. Injected embryos were screened for GFP and mCherry fluorescence under a Leica M165C stereomicroscope around 10-11hpf, when GFP reporter fluorescence first becomes detectable.

Selected embryos were immediately mounted in a phytagel tube (SIGMA, CAS Number: 71010-52-1) molded with Phaseview Teflon mold (1.5mm of diameter) and maintained in position with 0.4% low melting point agarose (Invitrogen UltraPure™ Low Melting Point Agarose). The tube containing the embryo was placed horizontally into the chamber containing 0.04% Tricaine in embryo medium (Sigma, CAS Number: 886-86-2). The tube was rotated under the microscope so that the embryo would face the objective.

Live imaging was performed approximately from 10.5-11hpf to 24hpf every 2.5min-3min, using a Phaseview Alpha^3^ light sheet apparatus, coupled with an Olympus BX43 microscope and using either a 20X/NA 0.5 Leica HCX APO objective or a 20X/NA 0.5 Olympus objective. Images were acquired using QtSPIM software (Phaseview), which controlled a Hamamatsu ORCA-Flash4.0 Digital sCMOS camera.

Room temperature was maintained at 24°C by air conditioning and the chamber temperature was further controlled by a BIOEMERGENCES-made thermostat. Medium level was maintained by a home-made perfusion system and an overflow to renew the medium.

### Movie analyses

#### Morphogenesis

Images were obtained and visualized with Arivis Vision4D software using re-oriented 3D stacks to allow similar optical section plane of analysis in different samples, cutting through the middle of the lens and the optic stalk at all time-steps. On one time-step per hour, measurements were performed on the re-oriented images: optic vesicle/optic cup length (at the widest), OV size increase (calculated by subtracting the length at the onset of furrow formation to the length at time t), optic stalk width, distance between the anterior optic cup and the lens, distance between the posterior optic cup and the lens, distance between the optic cup edges, position of the lens relative to anterior OV (=distance between center of the lens and anterior OV / (distance between center of the lens and anterior OV + distance between center of the lens and posterior OV) (see schemes on **Fig. 2** and **Fig. S4**).

#### Image stack treatments for cell tracking

Hyper-stacks used for tracking analyses were in 8-bit format. Pixel dimensions were 0.3 μm in x y, 1 μm in z, 39 t frames (2min30 each) and 420 and 360 z steps, respectively for surface fish and cavefish embryo. To improve image quality and allow more convenient tracking in MAMUT, several image treatments were necessary. Pixel intensity of all images within each stack were homogenized using contrast enhancement (0.3%), and 3D drift correction to improve image alignment was performed. Image stack were registered in the H5 format.

#### Cell tracking

To study cell behaviors, we tracked cell nuclei during evagination, between 11.5hpf and 13hpf (1h40, 40 movie frames) using the Fiji plugin MAMUT (Schindelin et al., 2012; Wolff et al., 2018) which allowed identification of nuclei at each t frame in the 3D. Because of the 3-fold increased voxel size compared to x and y, nuclei appeared distorted in the z plane. We preferentially –but not exclusively-tracked nuclei of high fluorescence intensity, which greatly facilitated non-ambiguous nuclei tracking. All nuclei tracks used for trajectory analyses were meticulously analysed and checked twice.

For trajectory analyses, the (x,y,z) cell coordinates were extracted using MAMUT and distances in 3D or 2D (x,y) between time points were calculated using the Pythagoras formula. We used x,y,z coordinates to calculate cumulative distance and absolute distance in space covered in 3D as well as instantaneous migration speeds (distance covered/150 sec). For the trajectory aspect, we used x,y coordinates to calculate instantaneous deviation angle at each time point using the Al-Kashi formula, valid in any triangle ABC, which relates the length of the sides using the cosine of one of the angles of the triangle. We calculated the value of the angle AB^AC in a triangle ABC, in which AB, BC and AC sides represent the distances covered by a nucleus between (t-t+1), (t+1-t+2) and (t-t+2), respectively. AB^AC=DEGRES(ACOS(((BC^2^)-(AB^2^)-(AC^2^))/-(ACxBC/2))).

To study proliferative activity, we tracked metaphases and anaphases manually and exhaustively in the whole brain /head of one SF and one CF embryo. To count mitotic events in OVs and presumptive lens without errors, each mitosis tracked and labelled in MAMUT was re-checked and allocated manually to structures or regions of interest (roi) (see **Fig. S6**). Results were expressed either as absolute cell counts, or normalized and expressed as densities to account for the difference of OV size between SF and CF (see **Fig. S6**). Two types of normalizations were applied, which lead to the same conclusion. First, the mitoses counts were normalized to the OV volumes, calculated on the movies using the plugin MZstack at 11.5, 12.5 and 13.5hpf and averaged (**Fig.3E**). Second, the mitoses counts were performed on maximum projections inside rois (regions of interest) of identical size, in the OVs or in the medial neural tube as a control (see **Fig. S6**). In the case of the OV roi, and because the optic vesicles are smaller in x,y but also in z (depth) in CF, a normalisation factor was applied. In SF, OV cell divisions were tracked along a z extent of 145, while in CF cell divisions were tracked on a z extent of 100. The normalisation factor was therefore x1.45 (**Fig. 3E**). For this proliferation analysis, statistical comparison could not be provided as we studied one SF and one CF sample.

### Statistics

Statistical significance and p-values were calculated using non-parametric Mann-Whitney U tests in R. No statistical method was used to predetermine sample size. The experiments were not randomized and the investigators were not blinded during image analyses.

## Acknowledgements

Work supported by an Equipe FRM grant (DEQ20150331745), UNADEV/AVIESAN and Retina France grants to SR. We thank Jean-Paul Concordet and Anne De Cian (Tacgene, Sorbonne Universités, Paris) for sharing Cas9 protein; Diane Denis, Krystel Saroul, Jocelyne Gaget, the Amagen personnel for advices and care of our *Astyanax* colony; Patrick Para for making custom tools for live microscopy; Adeline Boyreau, Adeline Rausch, Fanny Husson, Elena Kardash and Nadine Peyrieras (BioEmergence, Gif sur Yvette, France) for reagents, discussions and advices on live imaging, and for the use of the SPIM and image analysis tools; Cyprian Wozniak, Arthur Le Bris and Gaël Launay from PhaseView (Verrière-le-Buisson, France) for technical support and development of tools on the light-sheet microscope; Guillaume Plongeon for volume analyses; Martin Pepin (Sorbonne Université) for writing an essential macro in FIJI and Romain Le Bars at the I2BC Imaging Plateform facility for assistance on the FIJI software; Jean Yves Tinevez and the Image Analysis Hub at the Institut Pasteur (Paris) for help with tracking strategy.

## Supplemental Figures and Legends

**Supplemental Figure 1:**
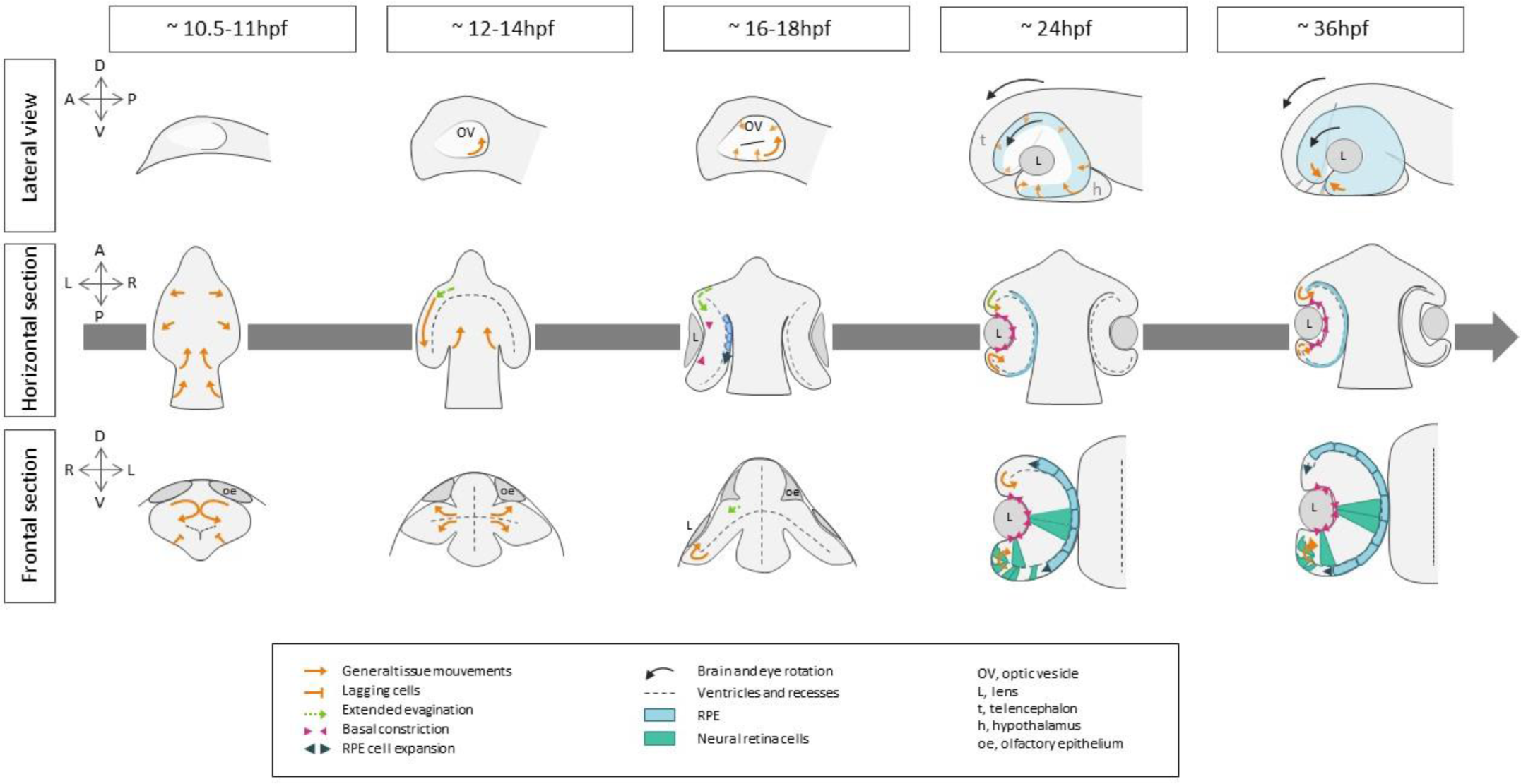
eye morphogenesis in fish. Schemes depicting the principal steps of eye morphogenesis in fish models, summarized from the available literature cited in Introduction. Stages and orientations are indicated. Orange arrows show general cell and tissue movements. Black arrows show the anterior-wise rotation of the eye and brain. Green arrows show the contribution of extended evagination. Pink arrowhead show cellular basal constriction. The blue color depicts the RPE cells, while the green color depicts retina neuroepithelium cells changing shape.

**Supplemental Figure 2:**
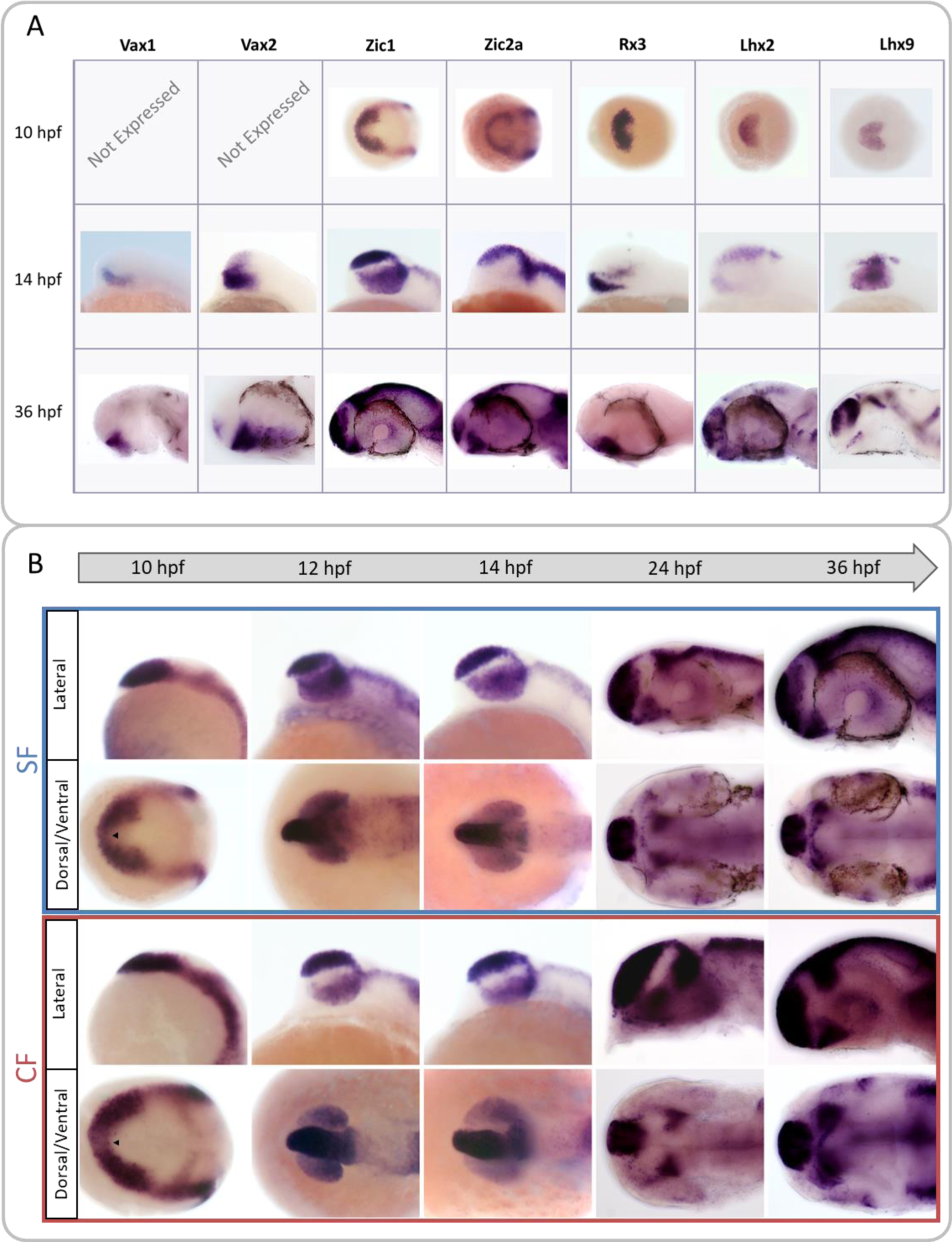
choosing a candidate gene for transgenesis. Chosen candidates were *Vax1*, *Vax2* (Take-uchi et al., 2003), *Zic1* (Hinaux et al., 2016; Maurus and Harris, 2009; Rohr et al., 1999; Tropepe et al., 2006), *Zic2a* (Sanek et al., 2009), *Rx3* (Deschet et al., 1999; Rembold et al., 2006; Stigloher et al., 2006), *Lhx2* and *Lhx9* (Pottin et al., 2011). (A) Mini-screen of candidate genes by *in situ* hybridization at different stages (10, 12, 14, 24 and 36hpf) of interest on surface fish and cavefish (not shown) embryos. Anterior is to the left. Dorsal views at 10hpf; lateral views at 14hpf and 36hpf. The eyes were dissected out for *Vax1* and *Lhx9* (as no eye expression was detected for either of them) to allow better visibility of the inner tissue. Among the 7 genes, 5 were expressed in the anterior neural plate at 10hpf while 2 were not: *Vax1* and *Vax2*, whose expressions were detectable from 12hpf only. Five of them were expressed at least partially in the OV per se (excluding ORR and optic stalk): *Vax2*, *Zic1*, *Rx3*, *Lhx9* and *Zic2a* (faintly). At 36hpf, only 4 of them were still expressed in the optic cup: *Zic2a* and *Zic1* (around the lens), *Lhx2* (faintly) and *Vax2* (in the ventral retina). Subtle differences between CF and SF expression patterns were observed (not shown), and only one candidate gene was consistently expressed in the eye from neural plate to 36hpf: *Zic1*. (B) Detailed analysis of *Zic1* expression pattern at 5 different stages in surface (SF) and cavefish (CF). Anterior is to the left, at 10, 12 and 14hpf, bottom pictures are taken in dorsal view; at 24 and 36hpf, bottom picture are taken in ventral views. Arrowheads indicate an indentation in the eyefield.

***Description of expression patterns:***

*Vax1* expression was detectable from 12hpf in the presumptive ORR (between the OVs) and additionally in the dorsal hypothalamus (according to brain axis (Puelles & Rubenstein, 2015), closest to the ORR) and quite faintly in the ventral telencephalon.

*Vax2* expression was very similar to *Vax1* both in terms of onset of expression and pattern, with the addition of the ventral quadrant of the eye. Although *Vax2* had a very interesting ventral pattern, we discarded it as a candidate for transgenesis for its expression onset was very late. Moreover, in *Vax2* enhancer trap zebrafish line (Kawakami Laboratory), the GFP fluorescence is only visible at 18hpf (personal observation, data not shown).

*Rx3* expression showed a typical eyefield expression pattern at 10hpf but progressively faded away during OV stages and was finally not expressed anymore at 24hpf. Conversely, an anterior and ventral expression in the presumptive hypothalamus was detectable from 12hpf and remained throughout the stages examined. At 36hpf, it was clear that only the dorsal half of the hypothalamus, closest to the ORR, was labelled. Due to the rapid fading of its OV expression, we did not consider *Rx3* as a valid candidate.

*Lhx2* and *Lhx9* were both already known to be expressed in the eyefield at neural plate stages in *Astyanax* (Pottin et al., 2011). *Lhx2* expression showed very dim expression, if any, in the OV at 12 and 14hpf but was expressed both in the prospective telencephalon and more faintly in the prospective hypothalamus. Later on at 36hpf, *Lhx2* was expressed strongly in the telencephalon and the olfactory epithelia; lighter expression was also visible in the ORR, hypothalamus and sometimes eyes. Additional expression in the pineal, optic tectum and in the hindbrain was also present.

*Lhx9* staining was strong in the OV at 12hpf (during evagination) and slightly lighter at 14hpf. Moreover dorsal and ventral lateral labelling at the border of the neural keel and the OV appeared, possibly prefiguring respectively the strong telencephalic staining visible at 24 and 36hpf and the hypothalamic cluster at the limit of the ORR already described in a previous publication (Alié et al., 2018). At these late stages, we could not detect *Lhx9* expression in the eye anymore. Salt and pepper staining was visible in the olfactory epithelia; a band of expression outlining the optic tectum and lateral discrete marks in the hindbrain were present. We did not choose *Lhx2* or *Lhx9* because of the rapid decay of their eye expression.

At 10hpf, *Zic2a* was expressed at the border of the neural plate and almost entirely surrounding the eyefield except for a medial posterior gap. Faint staining in the bilateral eyefield could also be seen on some embryos. At 12 and 14hpf, there was a strong *Zic2a* expression in the telencephalon and a faint staining in the eye or distal part of the eye could often be seen. Strong staining was generally visible throughout the dorsal-most brain. At 24hpf, *Zic2a* expression remained strong in the telencephalon and was also now strongly visible at the border of the eye, in the ORR or optic stalk but without reaching the midline. Faint staining in the eye remained. At 36hpf, the expression pattern was similar, with the ORR/optic stalk staining reaching much closer to the midline. The eye expression was now more focused around the lens, probably in the CMZ. Roof plate staining persisted throughout development. Because *Zic2a* was never strongly expressed in the eye, we did not favour it as a candidate for transgenesis.

*Zic1* was strongly expressed at 10hpf in the neural plate border and in the anterior neural plate, at the level of the eyefield. At 12 and 14hpf, *Zic1* expression was consistently found in the OV and between them (prospective ORR and optic stalk). A strong staining was also present throughout the telencephalon. More posteriorly, the roof plate of the midbrain and hindbrain was stained. The somites were also labelled. The pattern was very similar at 24hpf and 36hpf with a strong telencephalic expression and a milder ORR expression (mainly laterally and posterior to the optic recess)/optic stalk and eye staining (widely around the lens). Roof plate and somites expression remained. Even though its pattern of expression was complex and encompassed a region wider than the optic region of interest, *Zic1* was chosen for transgenesis due to its early and persistent expression throughout the eye and the ORR/optic stalk regions. Moreover, *Zic1* expression highlighted morphological differences between SF and CF. At 10hpf, in CF *Zic1* was expressed in narrower lateral bands in the eyefield, with a larger medial indentation. At 12hpf, *Zic1* pattern confirmed that the CF OV were shorter and “plumper”. At 36hpf the *Zic1*-expressing ORR was wider in cavefish.

**Supplemental Figure 3:**
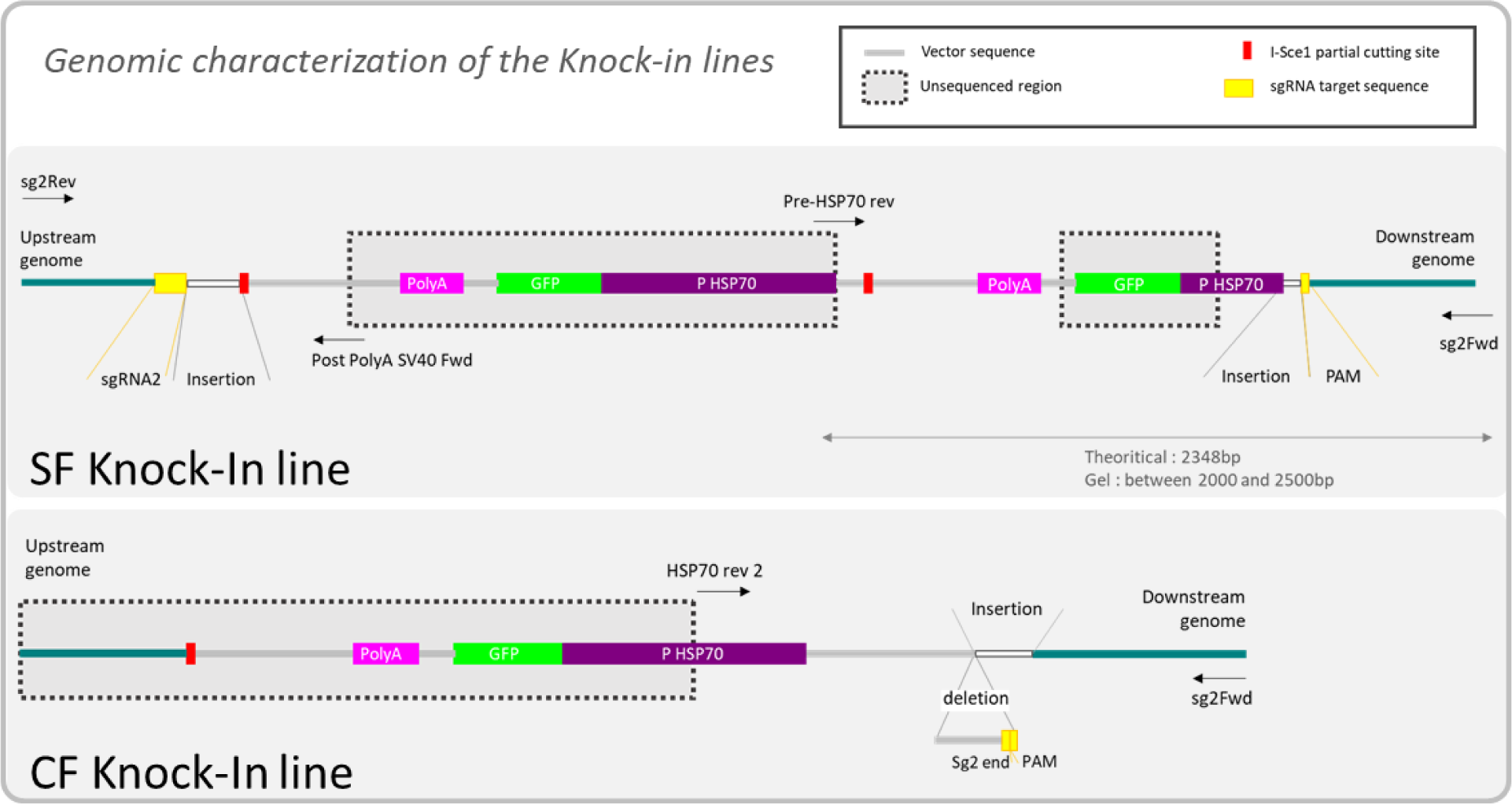
genomic characterization of the Knock-in lines. Knock-In insertions, based on partial sequencing. Dotted boxes indicate un-sequenced regions, leaving uncertainties. For example, in the surface fish line, there is at least a partial insertion of the repair construct, containing a truncated Hsp70 promoter and at least another insert in the same direction (but potentially several). Of note, the surrounding genomic region is very rich in T and A (GC content around 35%) with many repeats, making PCRs sometimes challenging. The data show that for both lines the transgenes are inserted at the correct targeted site.

**Supplemental Figure 4:**
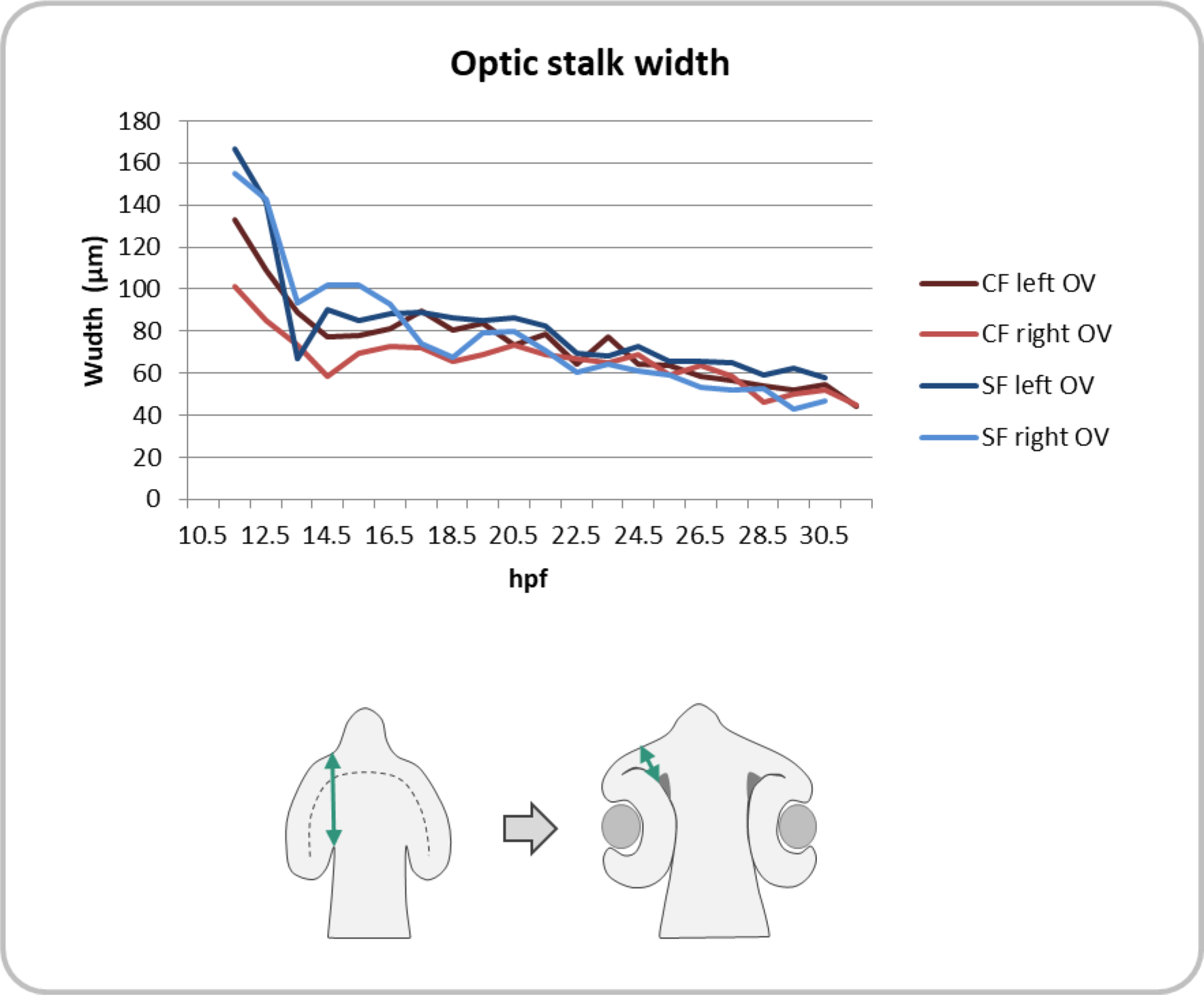
optic stalk width. The size of the optic stalk (in a wide meaning: the connection between the OV and the neural tube) is smaller in cavefish during early development due to the smaller size of the OV but rapidly becomes indistinguishable from the optic stalk of the surface fish.

**Supplemental Figure 5:**
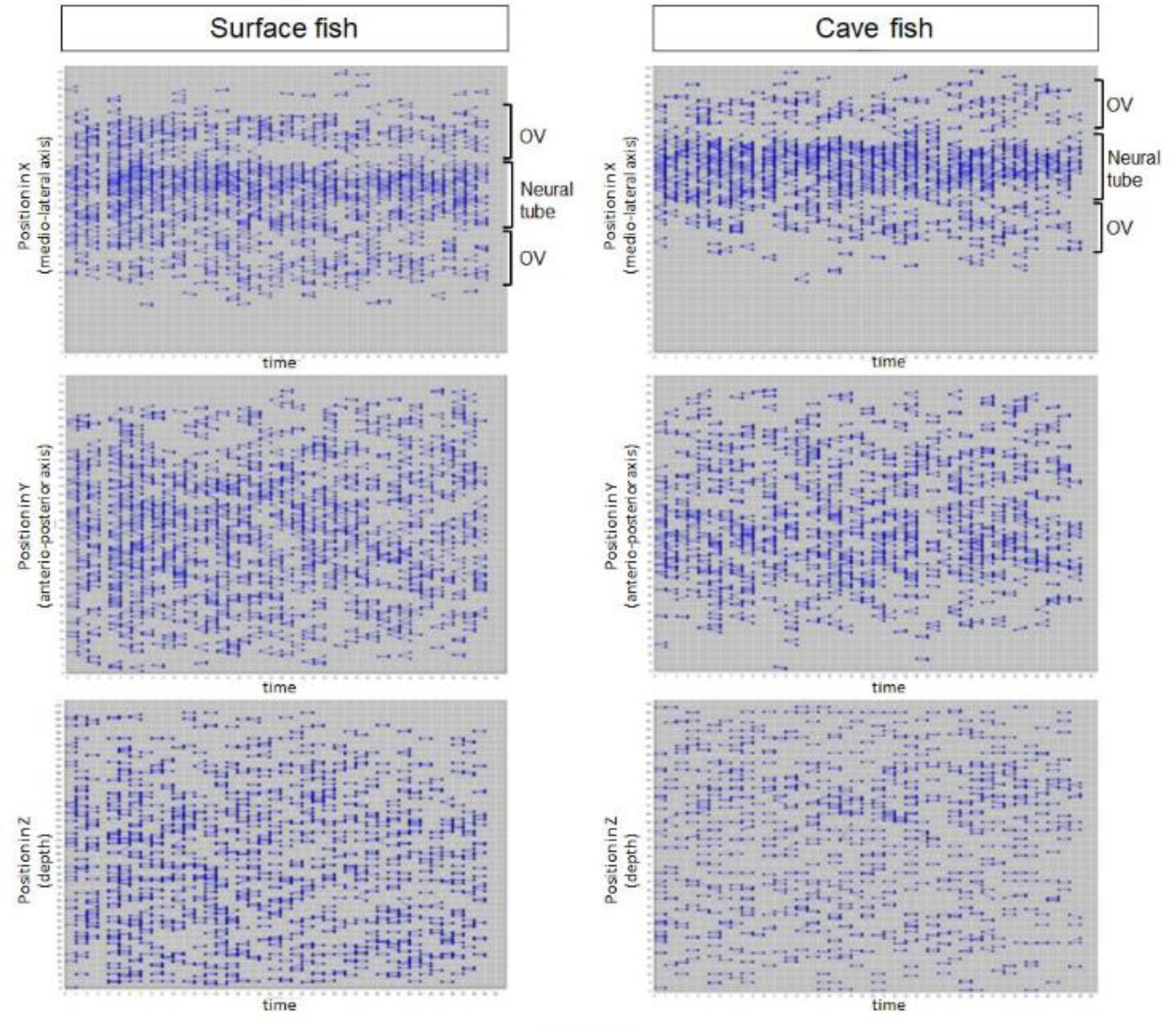
Distribution of mitoses in the head and as a function of time, between 11.5hpf and 13hpf, in SF and CF embryos. Plots showing the distribution of mitoses (schematically represented by a mother cell linked to daughter cells) in SF (left) and in CF (right). The 3 plots show the distribution of mitoses in X (medio-lateral axis), Y (antero-posterior axis) and Z (depth), as a function of time. Note the homogeneous repartition of divisions in the tissue, including in Z, suggesting that mitoses could be properly tracked, even in the depth of the tissue.

**Supplemental Figure 6:**
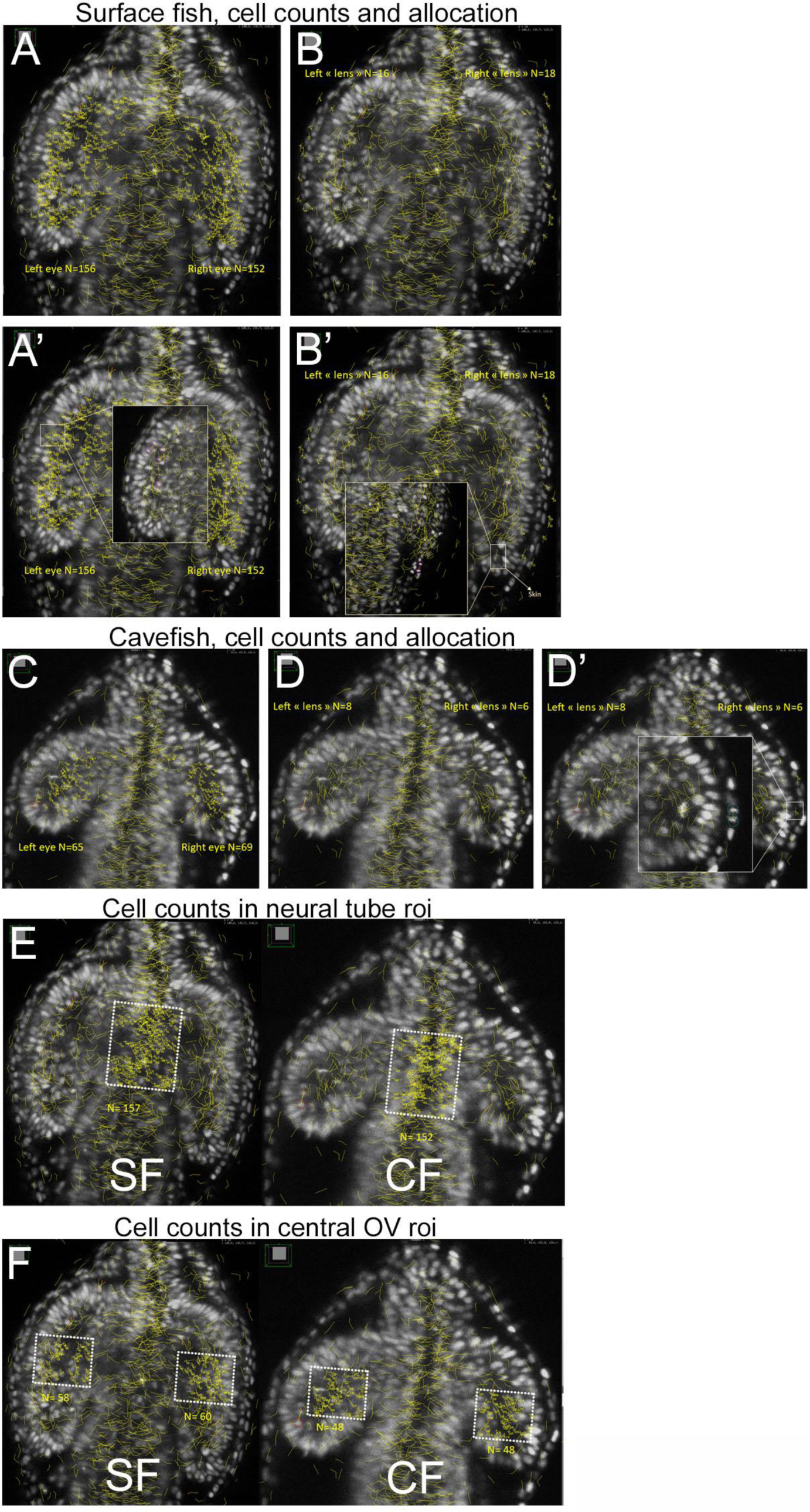
Counting mitoses, and normalization. (A-D’) Illustration of the method used to count mitoses in SF (A-B’) and CF (C-D’). To count mitoses in OVs and presumptive lens without errors, each mitosis tracked and labelled in MAMUT/Fiji was re-checked and segmented manually for proper allocation. Insets in A’B’D’ show examples of cells that appear like they belong to the OV region on the maximum projection, but that were attributed either to the OV, the skin or the lens after manual re-segmentation. (E) Illustration of mitosis counts in a medial neural tube roi of the same size in SF and CF, for estimation of mitotic density in the tissue. (F) Illustration of cell counts in OV roi of the same size in SF and CF for estimation of mitotic density in the tissue. Here, because the CF optic vesicles are smaller in XY but also in Z (depth), a normalisation factor was applied. In SF, OV cell divisions were tracked along a Z extent of 145, while in CF cell divisions were tracked on a Z extent of 100. The normalisation factor was therefore x1.45 (**Fig. 3E**).

**Supplemental Figure 7:**
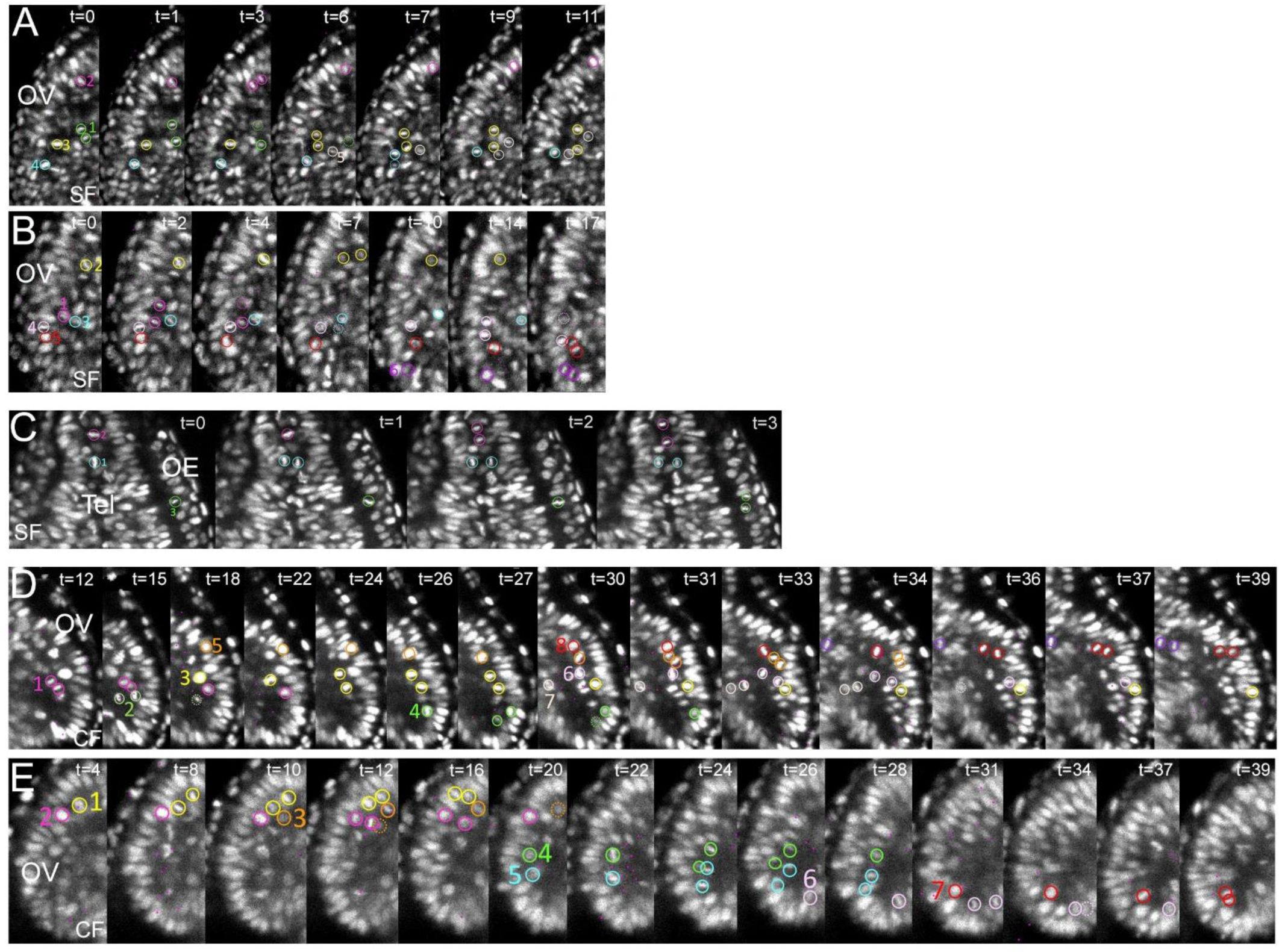
Additional examples of mitotic behaviors at high power magnification, with comments. Cell nuclei are labelled with colored circles; numbers indicate the order in which tracked cells will divide; daughter cells that migrate and that are lost in Z in the plane shown are indicated by dotted circles. (A,B) show the same cell divisions sequence as in Figure 3, in the SF OVs. <u>Comment for A:</u> The pink cell (#2) divides along the ventricle in the anterior OV and its daughter cell migrates and rapidly integrates in the neuroepithelium. The green and the beige cells (#1 and #5) divide in the proximal side of the ventricle and their daughter cells move towards the inner leaflet of the OV. So does the yellow cell (#3), although its initial position is more distal. Note the rotation/orientation behavior of the metaphasic plate of the blue cell, before dividing (#4). (C) shows cell divisions in the telencephalon (Tel) and the olfactory epithelium (OE) of SF. The pink and the blue cells (#1 and 2) divide in the telencephalon along the ventricular border, with orthogonally-oriented metaphasic plates. The green cell (#3) divides in the olfactory epithelium. (D,E) show cell divisions sequences in the CF OVs. D is the same sequence as in Figure 3. <u>Comments for D</u>: The pink (#1), yellow(#3), rose (#6) and red (#8) cells are representatives of all those cells that divide at the ventricle and then rapidly migrate to incorporate in the neuroepithelium. The orange cell (#5) follows a typical complete sequence: delamination from neuroepithelium, division at the ventricle, and reintegration of daughter cells back in the neuroepithelium. The kaki (#2) and the beige (#7) cells divide and populate the inner leaflet of the OV. The purple cell divides at the level of the optic recess region (ORR).

**Supplemental Figure 8:**
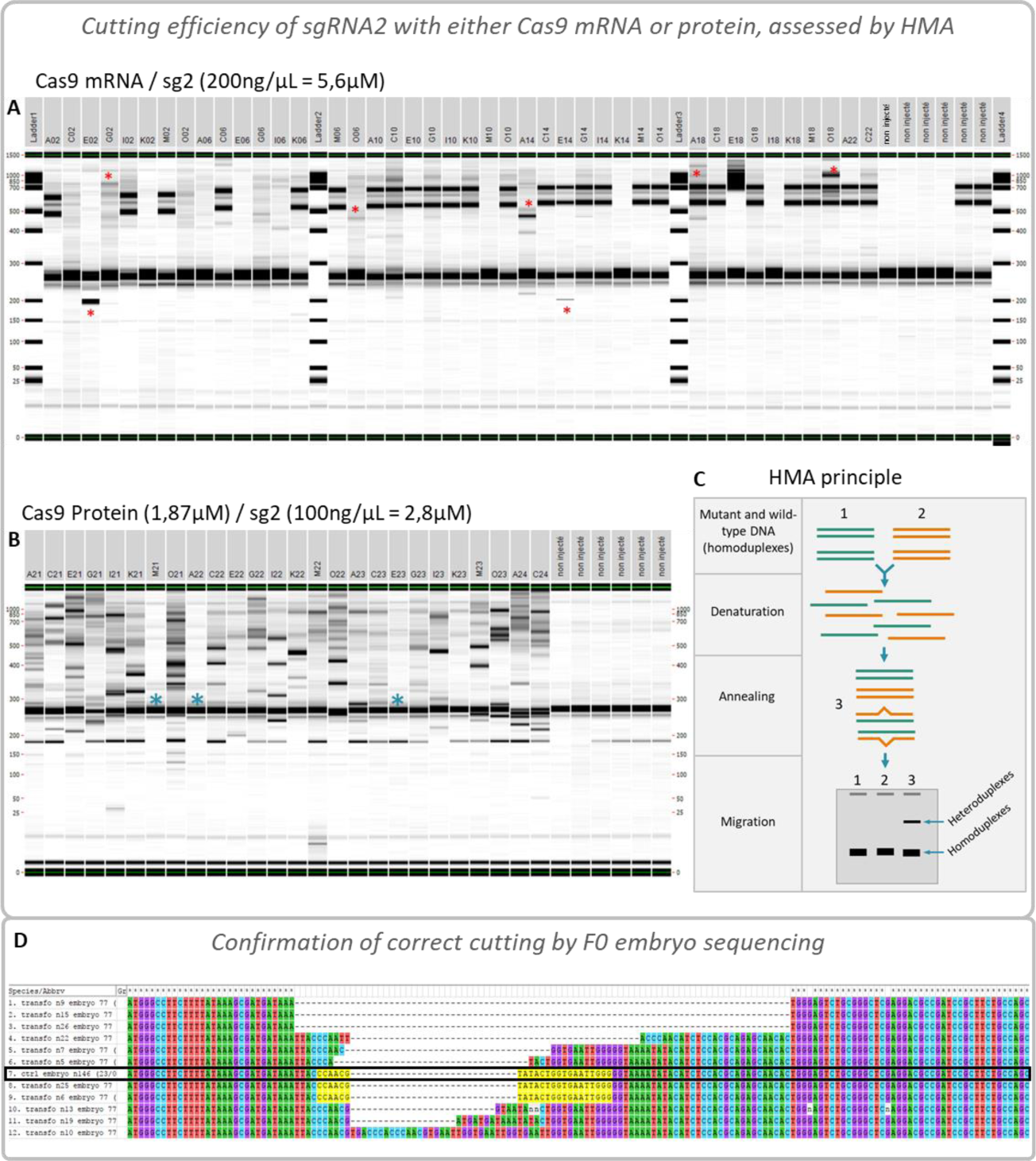
cutting efficiency of sgRNA 2. (A) Assessment of sgRNA 2 cutting efficiency when injected with Cas9 mRNA by heteroduplex mobility assay (HMA, explained in (C)). Each column is an individual F0 embryo. Embryos with strong additional bands are labelled with a red asterisk; additional light bands can be seen in several individuals, indicating cuts and imprecise repairs. Note that the 2 heavy bands seen on many embryos are also present in some of the un-injected controls (the 6 columns on the right) indicating a polymorphism in this region in the wild-type fish (not on the sgRNA target sequence). (B) Assessment of sgRNA 2 cutting efficiency when injected with Cas9 protein, note the strong presence of additional band compared to the 6 control embryos on the right. Embryos without any visible cuts are labelled with a blue asterisk. Additional bands are seen much more frequently and are much more important than with the Cas9 mRNA injection, probably indicating more frequent but also more precocious cut and repair events in the embryo, so that many cells share the same sequence. (C) Principle of the heteroduplex mobility assay: in an electrophoresis, heteroduplexes are slowed down compared to homoduplexes so that they form additional bands that can be seen even if the polymorphism is only a single substitution. In short, the DNA fragments are denatured and renatured to form heteroduplexes. An electrophoresis is then performed (here with a LabChip, PerkinElmer) to detect the presence of polymorphism. (D) Different cutting and repair events in a single injected embryo. A PCR was performed on one injected embryo (100ng/µL sgRNA2, Cas9mRNA) around the sgRNA2 target site and the product was cloned into pGEM-T Esay (Promega) and transformed into One shot TOP10 competent bacteria (Thermo Fischer). Plasmidic preparations from individual colonies were then sequenced. Various sequences were obtained, evidencing different cut and repair events in one single embryo. sgRNA2 target sequence is highlighted in yellow whenever intact. This F0 fish harbours both insertions and deletions around the cutting site of sgRNA2. A non-injected control fish sequence is included, outlined in black.

